# Transposable elements are important contributors to standing variation in gene expression in *Capsella grandiflora*

**DOI:** 10.1101/289173

**Authors:** Jasmina Uzunović, Emily B. Josephs, John R. Stinchcombe, Stephen I. Wright

## Abstract

Transposable elements (TEs) make up a significant portion of eukaryotic genomes and are important drivers of genome evolution. However, the extent to which TEs affect phenotypic variation on a genome-wide scale in comparison with other types of variants is still unclear. We characterised TE insertion polymorphisms and their effects on gene expression in 124 whole-genome sequences from a single population of *Capsella grandiflora*, and contrasted this with the effects of rare single nucleotide polymorphisms (SNPs). The frequency of insertions was negatively correlated with distance to genes, as well as density of conserved non-coding elements, suggesting that the negative effects of TEs on gene regulation are important in limiting their abundance. Rare TE variants strongly influence gene expression variation, predominantly through downregulation. In contrast, rare SNPs contribute equally to up- and down-regulation, but have a weaker effect than TEs. Taken together, these results imply that TEs are a significant contributor to gene expression variation and are more likely than rare SNPs to cause extreme changes in gene expression.

## INTRODUCTION

A key goal in evolutionary genetics is to understand the factors maintaining genetic variation in populations. With the advent of whole genome sequencing technologies, our ability to characterize the genetic basis of phenotypic variation has enabled new insights into this goal. The vast majority of studies examining the genetic basis of traits have focused on the influence of single nucleotide polymorphisms (SNPs), which in most populations are the most abundant class of variant and easiest to characterise using short-read whole genome sequencing (Atwell et al. 2010; Burke et al. 2014; Visscher et al. 2017). However, despite their numbers, SNPs are also likely to have relatively small individual effects compared with larger structural variants such as large insertions, duplications, deletions and rearrangement events, since they only affect the sequence of a single base pair of DNA. Consistent with the potential importance of structural variants, a recent study in humans suggested that, despite representing considerably less than 1% of called variants, structural variants appear to be causal at about 5% of expression quantitative trait loci (eQTLs), and they have larger effect sizes compared to SNPs (Chiang et al. 2017). Transposable elements (TEs), an important type of structural variation, have been shown to mediate large phenotypic changes in a number of cases (Daborn et al. 2002; Schlenke and Begun 2004; Butelli et al. 2012; Carrier et al. 2012; Van’t Hof et al. 2016). However, little is known about the genome-wide impact of TEs vs SNP variation on phenotypic variation.

Gene expression variation is a major contributor to phenotypic variation, leading to a growing interest in identifying genetic variants affecting expression (Brem and Kruglyak 2005; Massouras et al. 2012; Battle et al. 2014; Josephs et al. 2015; GTEx Consortium et al. 2017). There is increasing evidence for a major role of rare variants in the maintenance of expression variation, suggesting that a large fraction of expression variation may be maintained by a balance between new mutations introducing genetic variation and the action of selection on expression removing these variants (Zhao et al. 2016; Kremling et al. 2018; Hernandez et al. unpublished). However, the relative role of different classes of genetic variants, and the overall importance of rare vs. common variants in maintaining expression variation remains highly debated (Montgomery et al. 2011; Li et al. 2017). Transposable elements (TEs) may be a significant source of genetic variation for gene expression, as shown in genome-wide studies in several model organisms (Cridland et al. 2015; Quadrana et al. 2016; Stuart et al. 2016). In general, these studies have found that TE insertions are significantly associated with changes in nearby gene expression, with a predominantly dampening effect in *Drosophila* (Cridland et al. 2015), and a more balanced up- and down-regulating effect in *Arabidopsis* (Quadrana et al. 2016; Stuart et al. 2016). These studies suggest that TEs may contribute in important ways to heritable variation in expression in addition to SNPs.

A genome-wide survey of eQTL loci from a single large population of the self-incompatible outcrossing plant *Capsella grandiflora,* showed approximately a third of genes are associated with local eQTLs (those present within 5 kb of the gene), and SNPs associated with expression differences show evidence for being under purifying selection (Josephs et al. 2015). However, Josephs and colleagues examined only common SNPs (MAF > 5%), and the role of other classes of variants in driving expression variation remains unclear. Recent work in this species also shows evidence that TE insertions near genes targeted by siRNAs are subject to greater purifying selection, suggestive of the importance of TE silencing in generating expression variation (Horvath and Slotte 2017). *Capsella grandiflora* thus provides an excellent opportunity to explore the relative role of SNPs vs. transposable element activity in maintaining genetic variation in a natural population.

In this study, we examined TE polymorphism and its effects on expression variation using the same population studied by Josephs and colleagues (2015). We identified insertions across all individuals and explored the relationship between TE population frequency, rates of crossing over, and the density of coding sites and non-coding conserved sequences. We next tested the effect of TEs on gene expression variation using genome-wide expression data, and compared the relative contribution of rare TEs and SNPs to gene expression variation.

## RESULTS AND DISCUSSION

We combined high-confidence TE calls from 124 *C. grandiflora* plants using the RelocaTE2 pipeline (Chen et al. 2017). In total, we found 95,740 TEs in the population with a mean of 2468 insertions per individual and a median of 2675 insertions per individual (range of 497 to 3131). There are more TEs in *C. grandiflora* than in the related species *A. thaliana:* previous work found that in 216 *A. thaliana* accessions, there were 23,095 unique (non-reference) TE variants (with a mean of 1657 per accession) (Stuart et al. 2016). The higher abundance of TEs in the *C. grandiflora* population could reflect higher TE activity in outcrossers vs selfers leading to more low frequency insertions (Charlesworth and Langley 1986; Slotte et al. 2013).

The frequency spectrum of TEs in the population shows a major skew towards low-frequency insertions compared to 4-fold degenerate SNPs, with a strong enrichment of insertions present in only one individual (Figure 1A). The trend of mostly low-frequency insertions is consistent with observations from other organisms and suggests there is strong negative selection acting against TEs (Novick et al. 2011; Tian et al. 2012; Cridland et al. 2013; Quadrana et al. 2016; Stuart et al. 2016; Laricchia et al. 2017).

**Figure 1.**
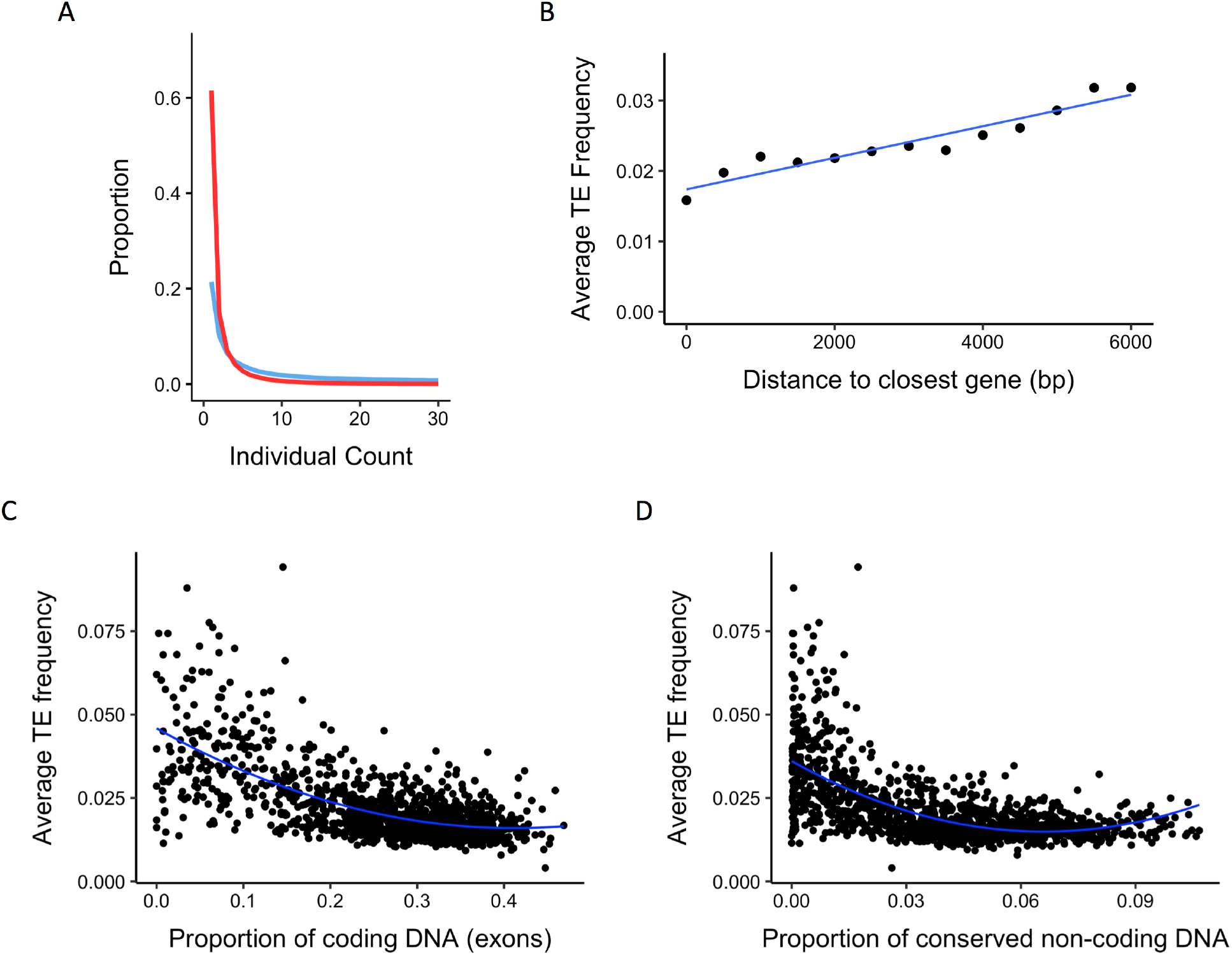
Patterns of TE insertion frequency from population data. (A) Frequency spectrum of TEs (red) and 4-fold SNPs (blue). (B) Relationship between TE frequency and distance to the closest gene. The first point represents the average frequency of all TEs which are within a protein-coding gene. Subsequent points average the frequency of all TEs in 500 bp windows from the closest gene. The blue line shows a linear model fit through the data. r = 0.94, linear coefficient has a p < 0.0001. C and D Relationship between average TE frequency and proportion of (C) coding DNA and (D) conserved non-coding sequence (CNS) in 100 kb windows across the genome. The blue curve shows a quadratic model fit. Spearman’s ρ = −0.562 in (C) and −0.619 in (D).

### Selection against gene disruption regulates TE numbers

There are two main hypotheses for why TEs are deleterious in the genome: (i) they can cause ectopic recombination between two non-homologous insertions or (ii) their insertion can disrupt gene function or expression. In the first hypothesis, TEs should be more deleterious in areas of high recombination rate, under the assumption that ectopic recombination rate mirrors the homologous crossover rate. We tested this hypothesis by comparing estimates of recombination rate in bins of 100 kb across the genome to the average TE population frequency of all insertions present in that bin (Figure S5). There is no significant correlation between TE frequency and recombination rate across the genome, suggesting ectopic recombination does not play a large role in outcrossing *C. grandiflora*, consistent with results from the highly selfing *A. thaliana* (Wright et al. 2003). Ectopic recombination is expected to be of greater importance in outcrossing species due to their higher heterozygosity; a TE is more likely to recombine non-homologously if it does not have a homologous partner to pair with. However, there is no evidence for a larger role of ectopic recombination in this outcrossing species than in a related selfing species. While finer-scale estimates of recombination might reveal a significant effect and it is possible that ectopic recombination rates are less correlated with rates of crossing-over than is often assumed, it appears that the previously observed lack of an effect of recombination on TE abundance in *A. thaliana* (Wright et al. 2003) cannot be attributed to its selfing mating system.

If TEs are deleterious because they disrupt gene function, insertions should be most deleterious, and consequently at lowest population frequency, when they are located in or near genes. There is a significant positive correlation between TE frequency and distance from gene in bins of 500 bp (Figure 1B; r = 0.94), suggesting that TEs closer to genes experience stronger purifying selection. This result is in line with a recent publication in *C. grandiflora* that utilized a pooled population sequencing analysis (Horvath and Slotte 2017).

Another way to test the gene disruption hypothesis is to plot the relationship between average TE frequency and coding site density in 100 kb windows across the genome. In testing this, we found a significant negative correlation, suggesting that TEs in regions of high gene density are kept at low population frequency (Figure 1C; Spearman’s ρ: −0.562). We also evaluated the relationship between TE frequency and the density of conserved non-coding sequences (CNSs), using CNSs conserved across nine Brassicaceae species, many of which show evidence of being regulatory elements (Haudry et al. 2013). There is also a strong negative correlation between TE frequency and CNS density, implying TEs face selective pressures due to negative effects of inserting into or near regulatory elements (Figure 1D; Spearman’s ρ: −0.619). There is an potential auto-correlation between coding site density and CNS density as regulatory elements will be found close to genes. To test whether the relationship of CNS density and TE frequency can be fully explained by the correlation between genes and CNSs, we built two quadratic regression models chosen because they showed the best fit to the data. In the first model, TE frequency is predicted only by proportion of coding sites while in the second, the proportion of CNSs is added as an additional predictor (the latter model is shown in Table 1; adjusted R^2^ = 0.502). Using an F-test on the two models showed the model with coding sites and CNSs explains significantly more variation than the model with coding sites alone (p < 0.0001). Similar analyses in *D. melanogaster* show that TEs are underrepresented in CNSs compared to the rest of the genome, however, there was no difference in population frequency of TEs in or outside of CNSs (Manee et al. 2018). Our results suggest that important TE effects on genes extend not only to direct gene disruption and methylation-mediated gene silencing, but also to disruption of regulatory sequence. The effect of TEs on CNSs could be due to direct disruption of enhancers, methylation of regulatory sequence caused by TE silencing, or spatial disruption of gene regulation caused by insertions between a gene and its regulatory region leading to aberrant expression.

**Table 1.** Quadratic regression model of average TE frequency.

| **Model: Average TE frequency ~ proportion of coding sequences (CSs) + (proportion of coding sequences)<sup>2</sup> + proportion of CNSs + (proportion of CNSs)<sup>2</sup>** |  |  |
| --- | --- | --- |
| **Coefficient** | **Estimate** | **P-value** |
| Proportion of coding sequences | -0.100 | < 0.0001 |
| (Proportion of coding sequences)<sup>2</sup> | 0.126 | < 0.0001 |
| Proportion of CNSs | -0.236 | < 0.0001 |
| (Proportion of CNSs)<sup>2</sup> | 1.399 | 0.000381 |
| **Adjusted R<sup>2</sup>** | 0.502 | < 0.0001 |

### TEs contribute significantly to gene expression variation

We have shown that TEs contribute to sequence variation, mostly as rare mutations that are under stronger purifying selection near genes and regulatory regions. Given this, we next investigated the extent to which TE insertions alter gene expression and contribute to standing genetic variation. One approach for testing for the effects of rare variants on expression is to ask whether individuals with extreme gene expression are often associated with TEs near their genes, or whether there is a greater burden of rare variants at the phenotypic extremes. Individual genes are sorted into expression rank bins, and then the total number of all local rare variants is plotted for all genes in all individuals. If rare variants cause a change in gene expression, either an increase or decrease, then this will result in a parabolic shape (Zhao et al. 2016).

To facilitate later comparison with SNP data, we used 109 individuals which passed TE, expression and SNP quality filters. Using all TEs with a MAF < 3% and within 500 bp of a gene, we found that individuals with more TEs near a gene are more likely to have extreme expression values than individuals without a nearby TE, and TEs appear to have a strong effect on decreasing gene expression and a weaker effect of increasing expression at the extremes (Figure 2A; R^2^ = 0.450). As a null control, we permuted mutation count and expression rank; as expected, permutations show no significant correlation, demonstrating the significance of our observed effect (Figure S6). This general result suggests that TEs do contribute to expression variation and that TE silencing and gene disruption effects may be more common than TE-mediated upregulation. This result is in contrast with a recent analysis in *A. thaliana*, where TE insertions driving increased expression appeared to be as prevalent as expression-decreasing effects (Quadrana et al. 2016, but see Stuart et al. 2016).

**Figure 2.**
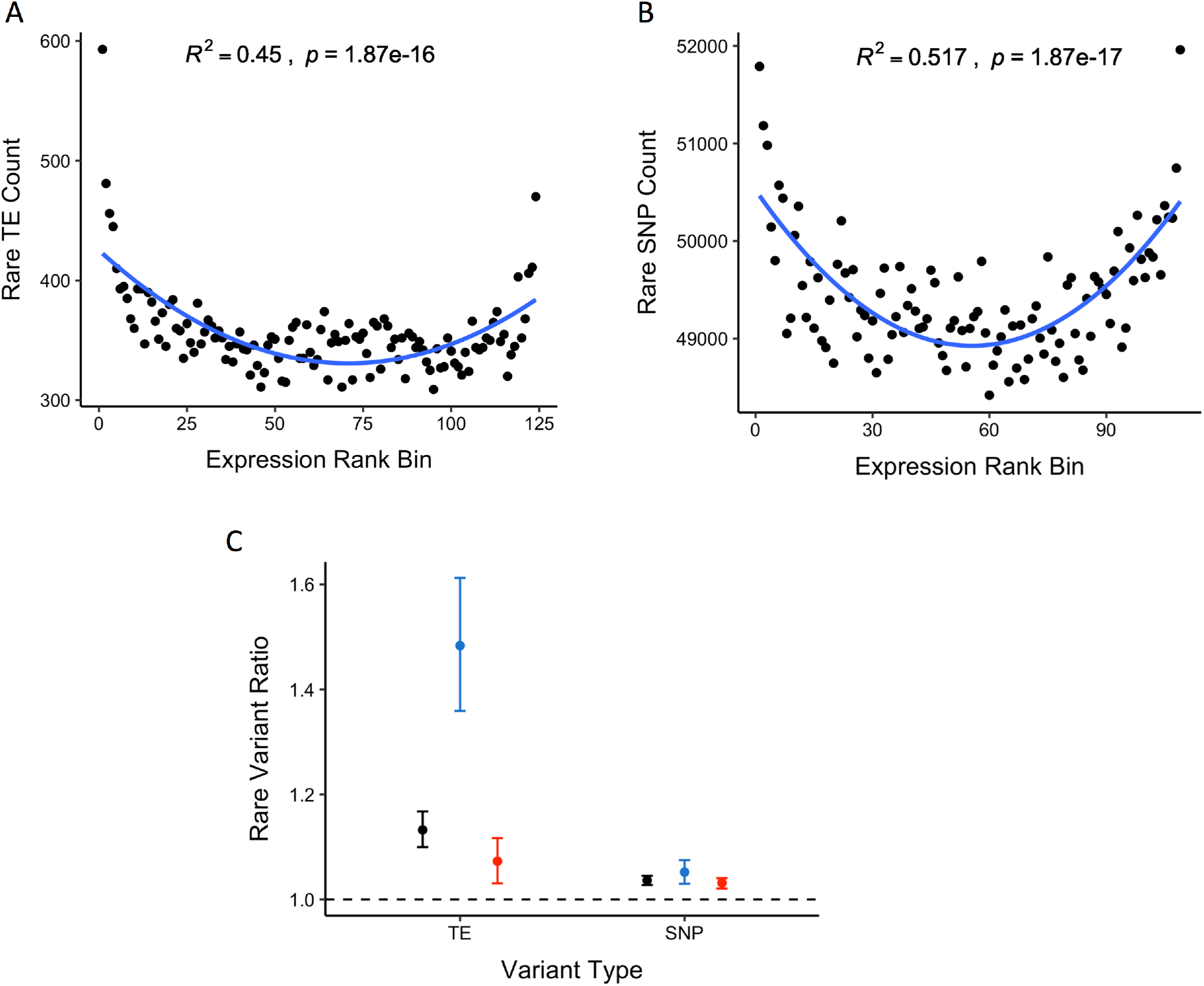
Relationship between (A) TE and (B) SNP count and gene expression rank bin for all rare variants (MAF < 3%) within 500 bp of a gene. C) Ratio of average number of rare variants in expression outliers relative to non-outliers for TEs and SNPs. ‘Upper’ refers to expression-increasing outliers, and ‘lower’ refers to expression-decreasing outliers. We ran 1000 permutations of the data to obtain normal distributions of the ratio statistic, and in all cases the observed statistic fell outside of the permuted distribution values.

While TEs appear to strongly downregulate gene expression, it is difficult from such an analysis to directly quantify the strength of the effect. We used a complementary statistical approach by calculating the ratio of the average number of TE insertions across genes in individuals that are expression outliers over the average number of TE insertions in non-expression outliers. The ratio value for TEs is 1.13, and expression outlier individuals have significantly more TEs than non-outlier individuals (Figure 2C; 95% quantile confidence interval: 1.10-1.17 from 1000 bootstraps). When considering only downregulated expression outliers, the ratio of TEs in outliers vs non-outliers is a much higher value of 1.48, consistent with the pattern observed in the burden of rare variants test (Figure 2C; 95% quantile confidence interval: 1.36-1.61 from 1000 bootstraps). These results suggest that the standing genetic variation contributed by TEs has a significant effect on disrupting gene expression, pushing individuals to extreme expression values and that TE insertions are more likely to decrease expression than increase it.

Breaking down the results by TE type reveals large differences in the direction and type of effect that different TE types have on gene expression. Long terminal repeat (LTR) elements, which are large retrotransposons, show the strongest effect on decreasing gene expression (R^2^ = 0.471 under the rare variant burden test) while other TE types tested show significant but less pronounced effects (Figure 3, Figure S7). LTRs are the most abundant TE type in most plant genomes, including *C. grandiflora* (Slotte et al. 2013; Bennetzen and Wang 2014), and often the most methylated (Ahmed et al. 2011). Methylated TEs, which have been silenced by the host genome to prevent their transcription and transposition, can have stronger effects on nearby gene expression, either through the spread of methylation past the TE or by making the chromatin less accessible to transcription factors and transcriptional machinery. Thus, the strong effect of LTRs on decreasing gene expression may be mediated by their methylation.

**Figure 3.**
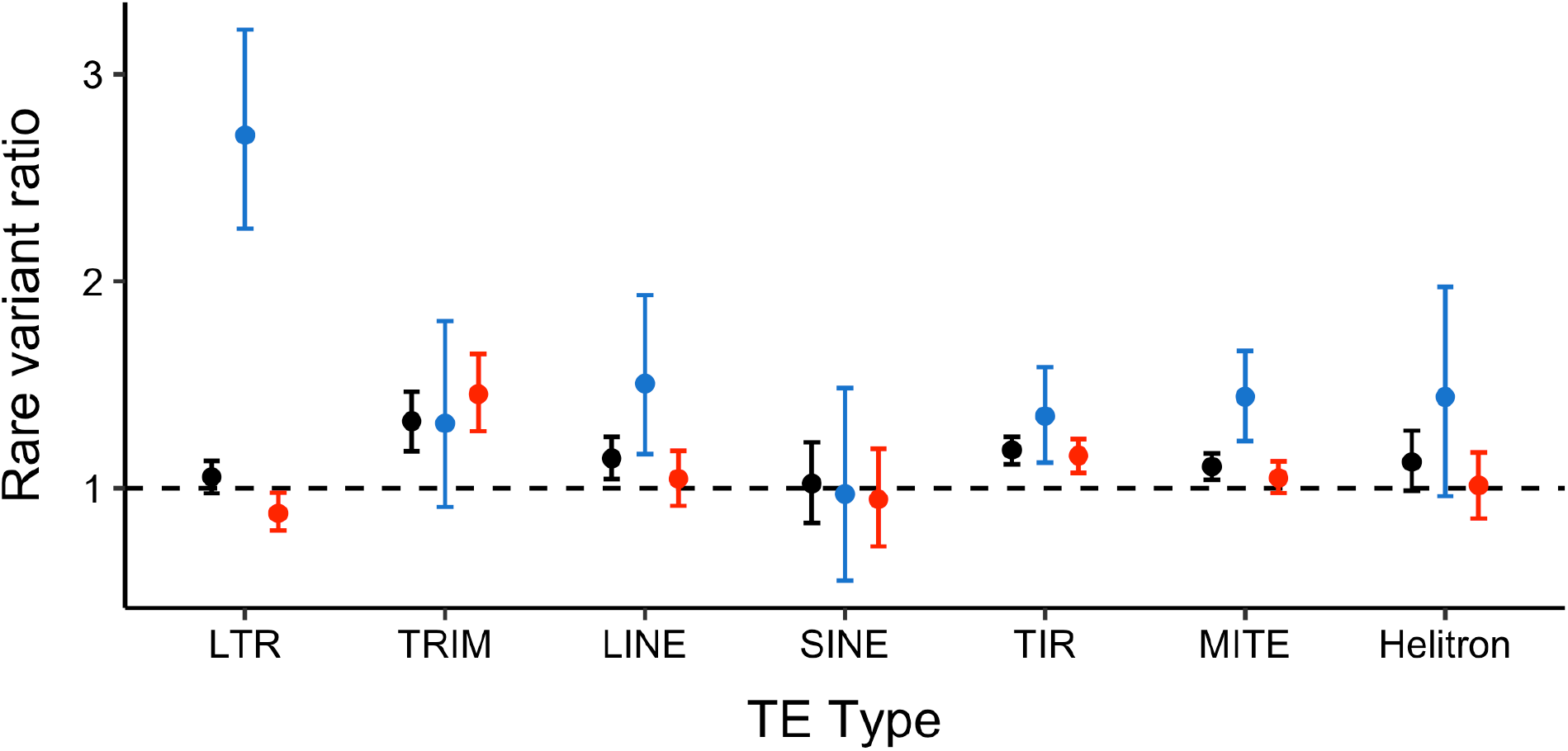
Ratio of average number of rare variants in expression outliers relative to non-outliers for different TE types. Black – all outliers; blue – expression-decreasing outliers; red – expression-increasing outliers.

### Genic insertions cause the strongest expression decrease

If TEs affect gene expression by disrupting gene regulatory regions, then there should be a strong effect caused by TEs upstream of the gene. However, the strongest correlation with extreme gene expression occurred when considering TEs inserted into genes (Figure S8; R^2^ = 0.524). Upstream TEs had a lower overall effect than inserted TEs; interestingly they were only significantly over-enriched in increased expression outliers, with no significant effect in decreased expression outliers (Figure 4). This is somewhat similar to observations in *A. thaliana*, where upstream insertions showed a more symmetrical effect on expression than insertions inside or downstream of genes (Stuart et al. 2016). The upstream effect was only significant for TEs within 100 bp of the transcription start site (TSS) (Figure 4, Figure S9), consistent with previous results suggesting that the greatest abundance of upstream conserved non-coding sites were 200 bp away from the TSS (Haudry et al. 2013). Similarly, upstream SNP eQTLs were most abundant within 1kb from the TSS (closer intervals were not tested) (Josephs et al. 2015). One possible explanation for the weaker and more symmetrical overall effect of insertions upstream of genes is that a major effect of TE insertions into regulatory sequence is an effect on expression noise, rather than a specific downregulation. Recent work on de novo mutational effects in regulatory regions in yeast (Metzger et al. 2016) suggest many more mutations have an effect on the variance in gene expression than on the expression mean. Similarly, TE effects in regulatory regions could be due to stochastic spreading of epigenetic silencing, driving a pattern of purifying selection on TEs associated with increased variance in average expression.

**Figure 4.**
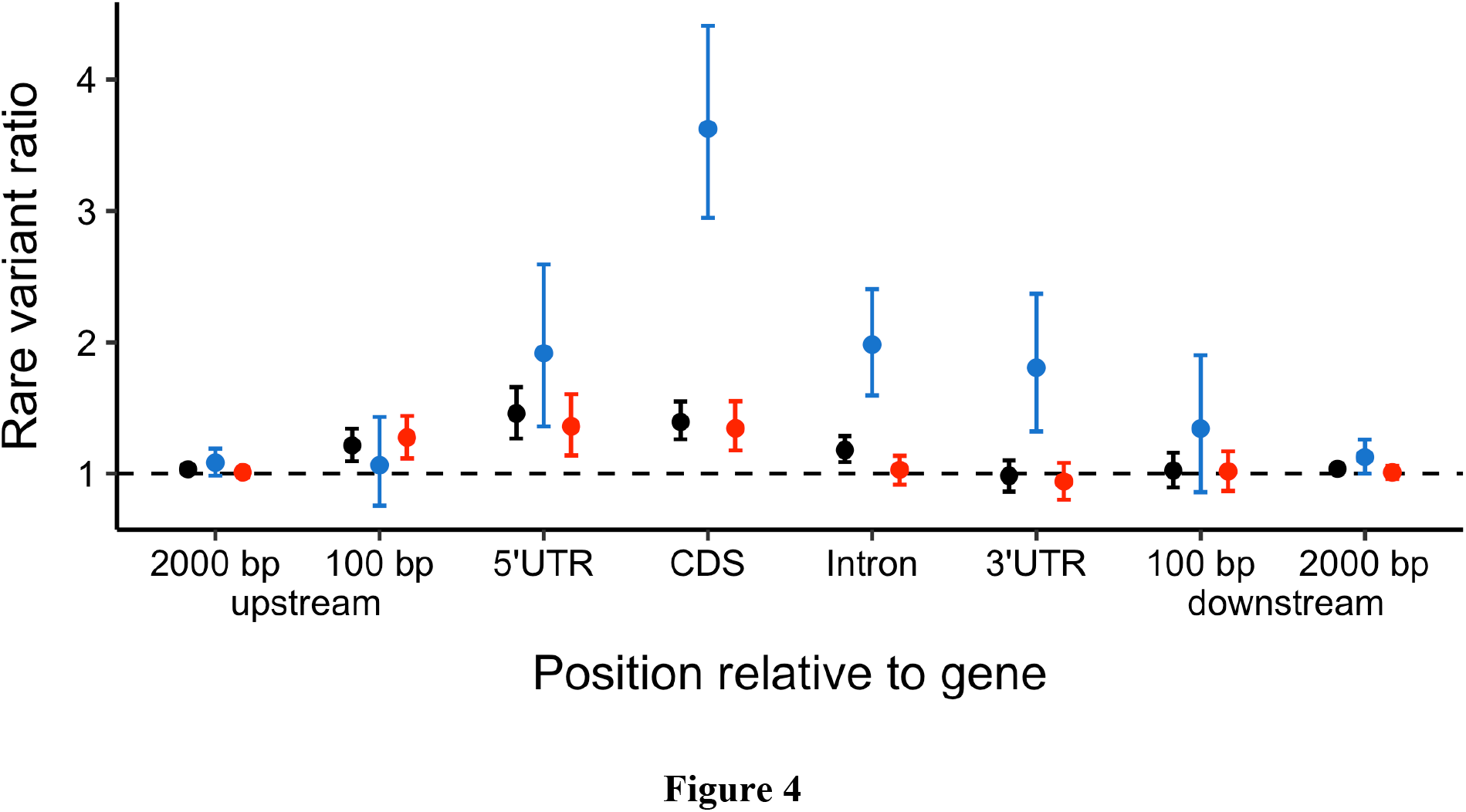
Ratio of average number of rare variants in expression outliers relative to non-outliers for TEs upstream of, in, and downstream of genes. The up- and downstream categories are non-overlapping. Black – all outliers; blue – expression-decreasing outliers; red – expression-increasing outliers.

Intragenic TEs appear to cause the strongest shift in expression, so we further categorised by the genic region of insertion. TEs in all genic regions were significantly over-enriched in low expression gene outliers, with the strongest effect from insertions into coding regions (Figure 4, Figure S10). In contrast, only TEs into the 5’UTR and coding sites were also significantly over-enriched in high expression gene outliers. The 5’UTR contains regulatory sites, so TE insertions may be contributing additional regulatory sites leading to increased transcription of the whole gene. TEs are known to be a large source of promoter and regulatory regions which can be co-opted by the host genome and contribute to the evolution of regulatory networks (Chuong et al. 2017; Hirsch and Springer 2017). The large effect of TEs in coding regions on decreasing gene expression could be due to faster decay of mRNAs with large mutations, for example by the nonsense-mediated decay (NMD) pathway (Garneau et al. 2007). However, the strong enrichment of TEs in low expression outliers when they insert into coding sites could be driven by a technical artifact where reads with TEs are dropped due to poor mapping (see Materials and Methods). When considering gene expression broken down into individual exons, there were no significant differences in enrichment of TEs in exons upstream of the insertion, downstream of the insertion and the exon containing the insertion (Figure S4). Therefore, there is not evidence for technical artifacts which could lead to misinterpretation of results. Furthermore, TEs appear to equally reduce expression of all exons, suggesting that TEs are not causing early transcription breaks. The impact of TEs in introns could be driven by targeted silencing of insertions in introns, as has recently been observed for L1 retrotransposons in humans (Liu et al. 2018).

### TE insertions show stronger enrichment in expression outliers than SNPs

Josephs and colleagues (Josephs et al. 2015) previously showed that a third of genes had an eQTL in this population of *C. grandiflora*, and that there is a significant negative correlation between eQTL MAF and effect size, suggesting that purifying selection is mostly responsible for maintaining genetic variation for gene expression. However, they tested SNPs with a MAF > 5%, and so the contribution of rare variants to gene expression variation is still unclear.

We explored the effect of rare SNPs with a MAF < 3% on gene expression. Overall, these variants are present in much larger number, and are equally likely to cause gene up- or down-regulation (Figure 2B; R^2^ = 0.517). This is in contrast with a recent study in maize, where SNP variants upstream of genes tended to show greater evidence of downregulation (Kremling et al. 2018). When comparing the ratio of variants in expression outliers vs non-outliers, SNPs had a smaller effect than TEs (1.04 vs 1.13, respectively; Figure 2C). As a class of mutation, TEs are 9% more over-represented in outliers than SNPs. In low expression outliers, there is a considerably greater gap between TE and SNP over-representation of 43% (1.48 vs 1.05 ratio values, respectively; Figure 2C). Both tests of the burden of rare variants on gene expression are genome-wide, made by summing across all genes and all individuals in the population. As such, the results highlight the global impact that both TEs and SNPs have on gene expression variation. The outlier test used a stringent cut-off of 4 standard deviations away from the mean to define outliers, so it may be that TEs are on average more likely to cause a large change in expression and are over-enriched in this test. SNPs, on the other hand, may be on average more likely to cause a small change in expression and are more important in explaining small variation in expression. Therefore, individual TEs may be strong candidates for having large effects on expression (large-effect QTL), but considering their numbers SNPs are also likely important, with smaller individual effects on average.

A major challenge when comparing SNPs to TEs is detection power. Calling non-reference TEs with short-read data is difficult, especially in outcrossing species where many insertions are heterozygous, leading to more false positives and false negatives than when calling SNPs (Ewing 2015). These power issues could lead to biases in our population frequency estimates. There may be many more rare TEs in the population which the pipeline is not detecting, however, in this case our results would be undervaluing the true effect sizes and are therefore conservative estimates of the impact of rare TEs. The outlier expression test controls for this problem because it calculates a ratio of TE numbers between two groups, both of which have equal detection power, allowing for the comparison of TE and SNP effect size independent of the ability to call these variants.

### Conclusions

We found that selection against disruption of gene function and regulation plays an important role in limiting TE copy number, but we see no evidence for selection against ectopic recombination in a population of *C. grandiflora*. TE insertions in or near genes strongly downregulate expression and to a smaller extent upregulate expression, with the effect differing by the type of TE and the genic region into which it has transposed. In comparison, SNPs have an overall weaker effect and are equally likely to up- and down-regulate expression. These results highlight the important contribution of TEs to standing genetic and phenotypic variation.

## MATERIALS AND METHODS

### Study organism

The self-incompatible, outcrossing plant *Capsella grandiflora*, is a model for using population genomic approaches to make inferences about natural selection due to its low population structure and large effective population size (N_e_ ≈ 600,000) (Foxe et al. 2009; Slotte et al. 2010; St.Onge et al. 2011). The species is a member of the Brassicaceae family and is native to Northern Greece and Southern Albania. There is evidence of prevalent purifying and positive selection on SNPs in *C. grandiflora* (Slotte et al. 2010; Williamson et al. 2014).

### Genomic data and expression

The DNA and RNA data have been previously described by (Josephs et al. 2015; Josephs et al. 2017). Briefly, plants were sampled from a single population located near Monodendri, Greece (population Cg-9 in St. Onge et al. 2011). Seeds were collected and grown in the University of Toronto glasshouses and random crosses were performed. The offspring of these crosses were grown in the University of Toronto growth chambers, where leaf tissue was collected and used for DNA and RNA extraction. Whole genome sequencing was done using 100 cycles of paired-end sequencing in a Hiseq 2000 with Truseq libraries (Illumina), resulting in 100 bp paired-end reads. RNA was sequenced at the Genome Quebec Innovation Centre in an Illumina HiSeq resulting in 100 bp paired-end or single-ended reads. For consistency, one read was randomly chosen among the paired end reads and they were subsequently treated as single ended. The reference genome used for mapping reads is *Capsella rubella*, a close relative of *C. grandiflora*, which diverged approximately 200,000 years ago (Foxe et al. 2009; Slotte et al. 2013).

RNA reads were mapped to an exon-only reference generated from the *C. rubella* reference genome (Slotte et al. 2013; Josephs et al. 2015) using Stampy 1.0.21 with default settings (Lunter and Goodson 2011). Expression level was measured with the HTSeq.scripts.count feature from HTSeq (Anders et al. 2015) and normalized for sequencing depth by dividing read counts by median read count of the entire sample. Genes with a median expression level below five reads per sample before normalization were removed from the analyses.

### Calling TEs

We built a new TE library to use with the detection pipeline, with a focus on TEs that show evidence of being recently active. Using pindel (Ye et al. 2009), we found large deletions (>150 bp) in 30 of the *C. grandiflora* short-read sequenced individuals relative to the *C. rubella* reference genome. These represent candidate TE insertions that occured in *C. rubella* since they in the reference genome at those locations but absent from those locations in the *C. grandiflora* sample. The PASTEClassifier pipeline (Hoede et al. 2014) allowed us to classify the sequences obtained from the *C. rubella* reference into TE order based on the hierarchical TE classification system (Wicker et al. 2007). Any sequences that remained unclassified were removed.

We used the RelocaTE2 (Chen et al. 2017) pipeline to detect both reference and non-reference TE insertions in 124 *C. grandiflora* individuals from our population. Briefly, RelocaTE2 is based on a combined split read and discordant read pairs mapping approach. All fastq files were subsampled down to 70X coverage to avoid elevated TE calls in individuals with higher coverage, and 23 individuals with below 70X coverage were removed. RelocaTE2 was run using bowtie2 as the TE aligner, using the custom-built TE library described above, and the *C. rubella* reference genome with default settings. We used bwa (Li and Durbin 2009) and samtools (Li et al. 2009) to align short reads to the reference genome, and then passed these alignments to RelocaTE2 to genotype TE insertions as heterozygous or homozygous. Heterozygous TE insertions will have reads aligning to the reference over the insertion site because one of the homeologs does not have a TE.

To create a population level table of TE insertions and their genotype (homozygous absent, heterozygous, or homozygous present), we merged all the insertions from the RelocaTE2 ‘high-confidence’ labelled files, which were insertions supported by at least one junction read from both sides. Any insertions called across individuals which were within 50 bp of one another and were of the same TE classification type were considered as the same insertion.

While RelocaTE2 can genotype insertions as homozygous or heterozygous, there is some evidence from the TE population frequency spectrum that these calls are not always accurate. When using allele counts (0, 1, or 2 for absent, heterozygous, homozygous) to make a frequency spectrum, there is a staggered pattern which goes away when examining the spectrum using individual count (0 or 1 for presence or absence; see Figure S1 and Figure 1). This suggests that homozygosity is being over-called. For all rare variant gene expression analyses (see below), we therefore used individual counts (the count of individuals with at least 1 TE insertion), effectively ignoring homozygous vs heterozygous calls, however our conclusions were qualitatively the same when performed with allele counts (Figure S2).

### TE validation

Validating RelocateTE2’s ability to find TEs is important, especially in outbred heterozygous population data where only half of the reads will support a heterozygous TE insertion. RelocaTE2 was previously tested on a rice strain of 20-fold genome coverage of 100 bp paired-end Illumina short reads aligned to the rice genome, and resulted in 93% specificity (the percent of calls which were true positives) (Chen et al. 2017). The *C. grandiflora* data should have a comparable level of specificity because it has higher coverage than the rice data previously tested; however the expected increased rate of heterozygous insertions may lead to a higher error rate. We used an independent bioinformatics test to estimate the true-positive rate (the proportion of new TEs called by RelocaTE2 that are actually TEs). We ran the SPAdes de novo assembler (Bankevich et al. 2012) on all TE calls on scaffold 1 for five individuals, chosen for their high coverage to increase the chance of success of the de novo assembler. The high coverage individuals should not bias the true-positive rate estimate because all individuals were subsampled to the same coverage (70X) for TE calling. We assembled all reads mapping to the region 1000 bp upstream and downstream of the TE insertion call and then aligned the de novo assembly contigs to the TE library using BLAST+ on default settings (Camacho et al. 2009). If we identified significant hits to the TE library, we considered the TE call to be a true positive.

The average true positive value across the five individuals was 80% for TEs labelled as high-confidence by RelocaTE2 (see Table S1 for breakdown by individual). The true positive value for non-high-confidence TEs was 60%. For subsequent analyses, we therefore focused on the high-confidence calls. Analyses considering both high-confidence and low-confidence calls led to similar results, but with lower significance, consistent with the greater noise contributed by the lower-confidence calls (Figure S3).

### Gene and genome annotation

TE insertions into genes were categorised as exonic, intronic, 3’UTR or 5’UTR based on the *C. rubella* genome annotation (Slotte et al. 2013). Similarly, 4-fold SNPs were identified for comparison with the TE site frequency spectrum. Conserved non-coding sites (CNSs) are determined through alignment of nine Brassicaceae species to the *C. rubella* genome as described in (Haudry et al. 2013; Williamson et al. 2014). Recombination rate data comes from fitting a third-order polynomial crossover events from an F2 genetic map derived from a cross between a *C. rubella* and *C. grandiflora* individual (Slotte et al. 2012; Slotte et al. 2013).

We ran quadratic multiple linear regression models predicting TE frequency from the proportion of coding sites alone, or both coding sites and CNSs using the R lm function (R Development Core Team 2018). We used the anova function to test whether the model with two predictor variables explained more variation than the simpler model.

### SNP dataset

Initial SNP calling and filtering was previously described (Josephs et al. 2015; Josephs et al. 2017). From the 124 individuals used in TE detection, 15 were dropped from further analyses due to an excess of NA SNP genotype calls (over 50,000 NA calls/individual). Any SNPs where more than nine individuals had an NA call were also dropped. The cutoffs were empirically determined as large outliers when plotting histograms of NA calls per individual and NA calls per SNP. All other NA calls were assumed to be homozygous for the reference allele; given the focus on rare variants, this assumption is unlikely to have a strong effect on results.

### Burden of rare variants test

The rare variant burden test used for determining the effect of rare variants (both TEs and SNPs) on gene expression was first described by Zhao and colleagues (Zhao et al. 2016). Briefly, the rare variant burden test estimates a genome-wide correlation between total rare variant count and gene expression relative to the rest of the population, testing whether individuals with more extreme expression values for a certain gene have more rare genetic variants located near that gene. Individuals are sorted into expression rank bins and the number of rare variants located in and near a gene is summed across all genes in the genome, for each expression rank bin. If there is a burden of rare variants, and having more of them pushes individuals to the extremes of expression (relative to the rest of the population), there will be a quadratic relationship between rare variant count and expression rank. Note that each gene is assessed independently, such that at individual genes, different individuals can end up in the tails of the distribution.

The rare variant burden plots were created using TEs and SNPs with a minor allele frequency (MAF) less than 3% from all 109 individuals. For all plots, we used TEs and SNPs which were within a gene or 500 bp upstream or downstream. We identified a small subset of genes where some individuals showed extreme numbers of rare SNPs, possibly reflecting alignment errors or paralogous sequences. Any genes where at least one individual had over 40 rare SNPs were removed (874 genes dropped in total).

For plots which considered only a single genic region (5’UTR, exon, intron, or 3’UTR), there may be some bias due to genes where UTRs were not annotated. Note that given the residual uncertainty of exact TE insertion locations (approx. 50 bp), this means that there will be some noise associated with annotation of insertion position.

### Expression outlier test

We developed an alternative method to test the effect of rare variants on gene expression which allows for a more direct comparison between the effect of TEs and SNPs. Again, we used all variants with a MAF < 3% from the 109 individuals which passed both TE and SNP filtering and which were located within a gene or 500 bp away from a gene.

The test involves determining individuals which are expression outliers, and comparing their number of rare variants with those of non-outlier individuals, repeated for each gene. Outliers were called as four standard deviations away from the mean, where mean and standard deviation were calculated using the expression values of individuals between the 1st and 3rd quartile. Estimating the mean and standard deviation from individuals between the 1st and 3rd quartiles reduces bias in these estimates caused by large expression value outliers and non-symmetrical distributions. As this method underestimates variation (relative to the total sample), an extreme cut-off of four standard deviations (SD) was used to call outliers. When testing without discrimination between upper and lower outliers, all genes were used except for 11, in which all individuals were within four SD of the mean (total of 18874 genes). When distinguishing between upper and lower outliers, we only considered genes where the mean was at least four SD above zero, to ensure it was possible to have lower outliers. After removing genes where all individuals were within four SD of the mean, there were 10290 genes tested for upper outliers and 14849 genes tested for lower outliers.

For every gene, we computed the number of rare variants per individual to control for differences in number of outliers and non-outliers among and between genes, and checked for a significant difference by running a Wilcoxon signed rank test using the R wilcox.test function for both TEs and SNPs (R Development Core Team 2018). To determine a per-variant effect size, we calculated the ratio of the mean number of rare variants per outlier individual to the mean number of rare variants per non-outlier individual across all genes. This ratio expresses how many more variants there are per outlier vs non-outlier. Taking the difference between the TE ratio and SNP ratio gives a quantitative measure of how much more likely the average TE is to cause an expression outlier vs the average SNP.

We ran 1000 bootstraps of the ratio using the R boot() package in order to calculate 95% quantile confidence intervals (Canty and Ripley 2017). 1000 permutations were also run to build a null distribution, where the outlier vs non-outlier label was switched with a 50% probability for every gene.

### Expression outlier by exon

We performed a slightly altered version of the expression outlier test to control for a possible technical artifact which could lead to the appearance of TE insertions in exons decreasing gene expression. RNA reads that include a TE may be dropped in filtering due to poor mapping, leading to an appearance of reduced overall transcript expression. To test this, we divided total gene expression into individual exon expression. For each gene with a TE insertion in at least one individual in the population, we created three categories: average expression of all exons upstream of the TE insertion, expression of the exon with the TE insertion, and average expression of all exons downstream of the TE insertion. If there was a technical artifact, and RNA reads with TEs were dropped due to poor mapping, we would expect that only exons with TEs had TEs associated with low expression outliers. Upstream and downstream exons wouldn’t show TE enrichment in low expression outliers, because their expression would not be affected by poor mapping.

## Data availability

All RNAseq and genomic data are available from the NCBI Sequence Read Archive (bioproject ID: PRJNA275635). File S1 contains all called TEs used in the analyses, their locations, and genotypes for each individual. File S2 contains all SNPs used in the analyses. All code used for analyses is available at https://github.com/jacau. All supplemental files and figures have been deposited at figshare.

## ACKNOWLEDGEMENTS

We thank Aneil Agrawal for key recommendations for statistical methods and Rob Ness for important suggestions on data validation. We thank Young Wha Lee, Niroshini Epitawalage, and Amanda Gorton for assistance collecting data. Thomas Bureau, Mathieu Blanchette, Daniel Schoen, Paul Harrison, Alan Moses, Adrian Platts, and Eef Harmsen contributed to the design and implementation of the Value-directed Evolutionary Genomics Initiative (VEGI). This work was supported by the Natural Science and Engineering Research Council of Canada (NSERC) (Discovery Grant to S.I.W. and J.R.S. and Canada Graduate Scholarships support to J.U.) and National Science Foundation (IOS-1523733 to E.B.J.).

## REFERENCES

Ahmed I, Sarazin A, Bowler C, Colot V, Quesneville H. 2011. Genome-wide evidence for local DNA methylation spreading from small RNA-targeted sequences in Arabidopsis. Nucleic Acids Res. 39:6919–6931.

Anders S, Pyl PT, Huber W. 2015. HTSeq—a Python framework to work with high-throughput sequencing data. Bioinformatics 31:166–169.

Atwell S, Huang YS, Vilhjálmsson BJ, Willems G, Horton M, Li Y, Meng D, Platt A, Tarone AM, Hu TT, et al. 2010. Genome-wide association study of 107 phenotypes in Arabidopsis thaliana inbred lines. Nature 465:627–631.

Bankevich A, Nurk S, Antipov D, Gurevich AA, Dvorkin M, Kulikov AS, Lesin VM, Nikolenko SI, Pham S, Prjibelski AD, et al. 2012. SPAdes: a new genome assembly algorithm and its applications to single-cell sequencing. J. Comput. Biol. 19:455–477.

Battle A, Mostafavi S, Zhu X, Potash JB, Weissman MM, McCormick C, Haudenschild CD, Beckman KB, Shi J, Mei R, et al. 2014. Characterizing the genetic basis of transcriptome diversity through RNA-sequencing of 922 individuals. Genome Res. 24:14–24.

Bennetzen JL, Wang H. 2014. The contributions of transposable elements to the structure, function, and evolution of plant genomes. Annu. Rev. Plant Biol. 65:505–530.

Brem RB, Kruglyak L. 2005. The landscape of genetic complexity across 5,700 gene expression traits in yeast. Proc. Natl. Acad. Sci. U. S. A. 102:1572–1577.

Burke MK, King EG, Shahrestani P, Rose MR, Long AD. 2014. Genome-wide association study of extreme longevity in Drosophila melanogaster. Genome Biol. Evol. 6:1–11.

Butelli E, Licciardello C, Zhang Y, Liu J, Mackay S, Bailey P, Reforgiato-Recupero G, Martin C. 2012. Retrotransposons control fruit-specific, cold-dependent accumulation of anthocyanins in blood oranges. Plant Cell 24:1242–1255.

Camacho C, Coulouris G, Avagyan V, Ma N, Papadopoulos J, Bealer K, Madden TL. 2009. BLAST+: architecture and applications. BMC Bioinformatics 10:421.

Canty, A, B. Ripley, boot: Bootstrap R (S-Plus) Functions. R package version 1.3-20. 2017.

Carrier G, Le Cunff L, Dereeper A, Legrand D, Sabot F, Bouchez O, Audeguin L, Boursiquot J-M, This P. 2012. Transposable elements are a major cause of somatic polymorphism in Vitis vinifera L. PLoS One 7:e32973.

Charlesworth B, Langley CH. 1986. The evolution of self-regulated transposition of transposable elements. Genetics 112:359–383.

Chen J, Wrightsman TR, Wessler SR, Stajich JE. 2017. RelocaTE2: a high resolution transposable element insertion site mapping tool for population resequencing. PeerJ 5:e2942.

Chiang C, Scott AJ, Davis JR, Tsang EK, Li X, Kim Y, Hadzic T, Damani FN, Ganel L, GTEx Consortium, et al. 2017. The impact of structural variation on human gene expression. Nat. Genet. 49:692–699.

Chuong EB, Elde NC, Feschotte C. 2017. Regulatory activities of transposable elements: from conflicts to benefits. Nat. Rev. Genet. 18:71–86.

Cridland JM, Macdonald SJ, Long AD, Thornton KR. 2013. Abundance and distribution of transposable elements in two Drosophila QTL mapping resources. Mol. Biol. Evol. 30:2311–2327.

Cridland JM, Thornton KR, Long AD. 2015. Gene expression variation in Drosophila melanogaster due to rare transposable element insertion alleles of large effect. Genetics 199:85–93.

Daborn PJ, Yen JL, Bogwitz MR, Le Goff G, Feil E, Jeffers S, Tijet N, Perry T, Heckel D, Batterham P, et al. 2002. A single p450 allele associated with insecticide resistance in Drosophila. Science 297:2253–2256.

Ewing AD. 2015. Transposable element detection from whole genome sequence data. Mob. DNA 6:24.

Foxe JP, Slotte T, Stahl EA, Neuffer B, Hurka H, Wright SI. 2009. Recent speciation associated with the evolution of selfing in Capsella. Proc. Natl. Acad. Sci. U. S. A. 106:5241–5245.

Garneau NL, Wilusz J, Wilusz CJ. 2007. The highways and byways of mRNA decay. Nat. Rev. Mol. Cell Biol. 8:113–126.

GTEx Consortium, Laboratory, Data Analysis &Coordinating Center (LDACC)—Analysis Working Group, Statistical Methods groups—Analysis Working Group, Enhancing GTEx (eGTEx) groups, NIH Common Fund, NIH/NCI, NIH/NHGRI, NIH/NIMH, NIH/NIDA, Biospecimen Collection Source Site—NDRI, et al.. 2017. Genetic effects on gene expression across human tissues. Nature 550:204–213.

Haudry A, Platts AE, Vello E, Hoen DR, Leclercq M, Williamson RJ, Forczek E, Joly-Lopez Z, Steffen JG, Hazzouri KM, et al. 2013. An atlas of over 90,000 conserved noncoding sequences provides insight into crucifer regulatory regions. Nat. Genet. 45:891–898.

Hernandez RD, Uricchio LH, Hartman K, Ye J, Dahl A, Zaitlen N, unpublished, https://www.biorxiv.org/content/early/2017/11/14/219238

Hirsch CD, Springer NM. 2017. Transposable element influences on gene expression in plants. Biochim. Biophys. Acta Gene Regul. Mech. 1860:157–165.

Hoede C, Arnoux S, Moisset M, Chaumier T, Inizan O, Jamilloux V, Quesneville H. 2014. PASTEC: an automatic transposable element classification tool. PLoS One 9:e91929.

Horvath R, Slotte T. 2017. The Role of Small RNA-Based Epigenetic Silencing for Purifying Selection on Transposable Elements in Capsella grandiflora. Genome Biol. Evol. 9:2911– 2920.

Josephs EB, Lee YW, Stinchcombe JR, Wright SI. 2015. Association mapping reveals the role of purifying selection in the maintenance of genomic variation in gene expression. Proc. Natl. Acad. Sci. U. S. A. 112:15390–15395.

Josephs EB, Wright SI, Stinchcombe JR, Schoen DJ. 2017. The relationship between selection, network connectivity, and regulatory variation within a population of Capsella grandiflora. Genome Biol. Evol. 9:1099–1109.

Kremling KAG, Chen S-Y, Su M-H, Lepak NK, Romay MC, Swarts KL, Lu F, Lorant A, Bradbury PJ, Buckler ES. 2018. Dysregulation of expression correlates with rare-allele burden and fitness loss in maize. Nature 555:520–523.

Laricchia KM, Zdraljevic S, Cook DE, Andersen EC. 2017. Natural Variation in the Distribution and Abundance of Transposable Elements Across the Caenorhabditis elegans Species. Mol. Biol. Evol. 34:2187–2202.

Li H, Durbin R. 2009. Fast and accurate short read alignment with Burrows-Wheeler transform. Bioinformatics 25:1754–1760.

Li H, Handsaker B, Wysoker A, Fennell T, Ruan J, Homer N, Marth G, Abecasis G, Durbin R, 1000 Genome Project Data Processing Subgroup. 2009. The Sequence Alignment/Map format and SAMtools. Bioinformatics 25:2078–2079.

Liu N, Lee CH, Swigut T, Grow E, Gu B, Bassik MC, Wysocka J. 2018. Selective silencing of euchromatic L1s revealed by genome-wide screens for L1 regulators. Nature 553:228–232.

Li X, Kim Y, Tsang EK, Davis JR, Damani FN, Chiang C, Hess GT, Zappala Z, Strober BJ, Scott AJ, et al. 2017. The impact of rare variation on gene expression across tissues. Nature 550:239–243.

Lunter G, Goodson M. 2011. Stampy: a statistical algorithm for sensitive and fast mapping of Illumina sequence reads. Genome Res. 21:936–939.

Manee MM, Jackson J, Bergman CM. 2018. Conserved Noncoding Elements Influence the Transposable Element Landscape in Drosophila. Genome Biol. Evol. 10:1533–1545.

Massouras A, Waszak SM, Albarca-Aguilera M, Hens K, Holcombe W, Ayroles JF, Dermitzakis ET, Stone EA, Jensen JD, Mackay TFC, et al. 2012. Genomic Variation and Its Impact on Gene Expression in Drosophila melanogaster. PLoS Genet. 8:e1003055.

Metzger BPH, Duveau F, Yuan DC, Tryban S, Yang B, Wittkopp PJ. 2016. Contrasting Frequencies and Effects of cis- and trans-Regulatory Mutations Affecting Gene Expression. Mol. Biol. Evol. 33:1131–1146.

Montgomery SB, Lappalainen T, Gutierrez-Arcelus M, Dermitzakis ET. 2011. Rare and common regulatory variation in population-scale sequenced human genomes. PLoS Genet. 7:e1002144.

Novick PA, Smith JD, Floumanhaft M, Ray DA, Stéphane B. 2011. The evolution and diversity of DNA transposons in the genome of the lizard Anolis carolinensis. Genome Biol. Evol. 3:1–14.

Quadrana L, Bortolini Silveira A, Mayhew GF, LeBlanc C, Martienssen RA, Jeddeloh JA, Colot V. 2016. The Arabidopsis thaliana mobilome and its impact at the species level. eLife 5:e15716

R Development Core Team, 2008 R: A Language and Environment for Statistical Computing. R Foundation for Statistical Computing, Vienna. http://www.R-project.org.

Schlenke TA, Begun DJ. 2004. Strong selective sweep associated with a transposon insertion in Drosophila simulans. Proc. Natl. Acad. Sci. U. S. A. 101:1626–1631.

Slotte T, Foxe JP, Hazzouri KM, Wright SI. 2010. Genome-wide evidence for efficient positive and purifying selection in Capsella grandiflora, a plant species with a large effective population size. Mol. Biol. Evol. 27:1813–1821.

Slotte T, Hazzouri KM, Ågren JA, Koenig D, Maumus F, Guo Y-L, Steige K, Platts AE, Escobar JS, Newman LK, et al. 2013. The Capsella rubella genome and the genomic consequences of rapid mating system evolution. Nat. Genet. 45:831–835.

Slotte T, Hazzouri KM, Stern D, Andolfatto P, Wright SI. 2012. Genetic architecture and adaptive significance of the selfing syndrome in capsella. Evolution 66:1360–1374.

St. Onge KR, Källman T, Slotte T, Lascoux M, Palmé AE. 2011. Contrasting demographic history and population structure in Capsella rubella and Capsella grandiflora, two closely related species with different mating systems. Mol. Ecol. 20:3306–3320.

Stuart T, Eichten SR, Cahn J, Karpievitch YV, Borevitz JO, Lister R. 2016. Population scale mapping of transposable element diversity reveals links to gene regulation and epigenomic variation. eLife 5:e20777

Tian Z, Zhao M, She M, Du J, Cannon SB, Liu X, Xu X, Qi X, Li MW, Lam HM, et al. 2012. Genome-wide characterization of nonreference transposons reveals evolutionary propensities of transposons in soybean. Plant Cell 24:4422–4436.

Van’t Hof AE, Campagne P, Rigden DJ, Yung CJ, Lingley J, Quail MA, Hall N, Darby AC, Saccheri IJ. 2016. The industrial melanism mutation in British peppered moths is a transposable element. Nature 534:102–105.

Visscher PM, Wray NR, Zhang Q, Sklar P, McCarthy MI, Brown MA, Yang J. 2017. 10 Years of GWAS Discovery: Biology, Function, and Translation. Am. J. Hum. Genet. 101:5–22.

Wicker T, Sabot F, Hua-Van A, Bennetzen JL, Capy P, Chalhoub B, Flavell A, Leroy P, Morgante M, Panaud O, et al. 2007. A unified classification system for eukaryotic transposable elements. Nat. Rev. Genet. 8:973–982.

Williamson RJ, Josephs EB, Platts AE, Hazzouri KM, Haudry A, Blanchette M, Wright SI. 2014. Evidence for widespread positive and negative selection in coding and conserved noncoding regions of Capsella grandiflora. PLoS Genet. 10:e1004622.

Wright SI, Agrawal N, Bureau TE. 2003. Effects of recombination rate and gene density on transposable element distributions in Arabidopsis thaliana. Genome Res. 13:1897–1903.

Ye K, Schulz MH, Long Q, Apweiler R, Ning Z. 2009. Pindel: a pattern growth approach to detect break points of large deletions and medium sized insertions from paired-end short reads. Bioinformatics 25:2865–2871.

Zhao J, Akinsanmi I, Arafat D, Cradick TJ, Lee CM, Banskota S, Marigorta UM, Bao G, Gibson G. 2016. A Burden of Rare Variants Associated with Extremes of Gene Expression in Human Peripheral Blood. Am. J. Hum. Genet. 98:299–309.

